# Reduced spread of nodes in spatial network models improves topology associated with increased computational capabilities

**DOI:** 10.1101/2024.10.09.617483

**Authors:** Nicholas Christiansen, Ioanna Sandvig, Axel Sandvig

**Affiliations:** Department of Neuromedicine and Movement Science, Faculty of Medicine and Health Sciences, Norwegian University of Science and Technology (NTNU), Trondheim, Norway; Department of Clinical Neurosciences, Umeå University Hospital, Umeå, Sweden; Department of Community Medicine and Rehabilitation, Umeå University, Umeå, Sweden

**Keywords:** neural architectures, communication efficiency, wiring, neuroplasticity, spatial models

## Abstract

Biological neural networks are characterized by short average path lengths, high clustering, and modular and hierarchical architectures. These complex network topologies strike a balance between local specialization and global synchronization via long-range connections, resulting in highly efficient communication. Efficient network organization and information processing are directly affected after network perturbation, e.g., as a result of trauma or neurodegenerative disease. Here, we used a spatial network model combining changes to spatial constraints on neuron placement with different wiring probabilities to investigate the effects of wiring cost principles on network complexity for different spatial conformations. By combining different wiring probabilities with varying levels of spatial clustering, we aimed to understand how alterations to mechanisms such as the neurons ability to group together, leading to different levels of clustering, in combination with altered axon outgrowth and branching, change network topology. Through targeted pruning of the networks we further aimed to understand how these topologies changed due to perturbations, resulting in a breakdown of function. We found that both long-range and intermediate wiring probabilities only conform to small-world architectures for neurons in dense spatial clusters due to a decrease in wiring cost within clusters. Furthermore, both small-worldness and modularity were reduced in systems with long-range connections caused by a reduction in network clustering. The presence of long-range connections were found to improve global information transmission, especially in networks with strong spatial clustering. In addition, these connections were shown to have a protective effect on the networks during targeted pruning, maintaining global connectivity even with high levels of pruning. Combining spatial clustering with wiring cost principles allows for novel insights into mechanisms underlying adaptive or maladaptive network alterations, explaining how specific interactions may lead to observed changes in networks. Our findings corroborate previous work showing that both wiring probability and spatial distributions play a key role in neural network development and response to perturbations, and increase our understanding of how maladaptive responses may lead to increased strain on local circuitry while negatively impacting global communication.

**Author summary:** Biological neural networks organize into networks capable of performing a variety of complex tasks in a cost-efficient manner. In networks where neurons are subject to neuropathological disease or perturbations, the mechanisms leading to these networks are disrupted, resulting in changes to the computational capabilities of the networks which can be challenging to elucidate. In this study, we combine spatial network models with different wiring costs used to establish connections between nodes of the network to gain an understanding of how changes to the wiring cost and spatial distribution of nodes in a network impacts the resulting topologies. We find that irrespective of wiring cost, sufficient spatial clustering is required for the emergence of network capabilities related to onset and maintenance of activity found in biological neural networks. Furthermore, we find that a balance between local connectivity and long-range connections improve outcomes when networks are subject to targeted perturbations of these connections. To summarize, spatial models prove an efficient method for understanding how changes related to wiring cost and targeted deletion of connections may change topologies of networks, allowing us to understand how specific changes to networks may arise and manifest in in vitro and in vivo cases.

## Introduction

An intrinsic property of biological neurons during normal development is their ability to self-organize into functional complex network architectures. These networks are formed through the establishment of connections between axons and dendrites, as neurites develop due to intrinsic growth programs, external guidance cues and growth factors, and an interplay between activity-dependent and activity-independent plasticity mechanisms (Onesto et al., 2021; Okujeni et al., 2017). The resulting networks balance metabolic consumption with the cost of long-range connections, while allowing for optimal capacity of information transfer (Laughlin, 2003; Beggs, 2008; Bullmore and Sporns, 2012). These organizational principles confer the capability of larger dynamical changes in the global activity necessary for the complex computations performed by the brain, as well as having a protective effect from perturbations to the network (Yamamoto et al., 2018; Kaiser, 2010; Pan and Sinha, 2007).

The study of these complex networks at different scales using concepts from network science allows for new insight into how network structure modulates activity (Rubinov and Sporns, 2010; Khambhati et al., 2018; Massobrio et al., 2015; Okujeni et al., 2017; Okujeni and Egert, 2019a). This has revealed several topological hallmarks of biological neural networks present across multiple scales, including their affinity for organizing into structures defined by a high degree of small-worldness, a hierarchical and modular organization, and a lack of characteristic scale (i.e. scale-free degree distributions) (Kaiser, 2010; Betzel et al., 2017; Goulas et al., 2019; Antonello et al., 2022). In neurodegenerative diseases such as amyotrophic lateral sclerosis (ALS), the mechanisms leading to these hallmarks of healthy networks are disrupted as a result of progressing pathology, with an increasing amount of literature pointing to alterations to genes related to cell adhesion and neurite outgrowth (Perkins et al., 2021; Kollstrøm et al., 2025a,b; Zimyanin et al., 2026). Specifically, in human ALS patient derived motor neurons we have previously demonstrated that diseased networks show aberrant axon branching and decreased axon outgrowth lengths. Ultimately, network function suffers, as maladaptive network responses to evolving pathology lead to changes in activity, increasing strain and placing a higher metabolic cost on centralized neurons (Fiskum et al., 2026).

Capturing these maladaptive responses in network models can be challenging. Most of the widely adopted models disregard effects from spatial constraints on network formation, relying on randomly assigning connections, rewiring of lattices or preferential attachment based on node degree, necessitating the need for alternative null models (Erdos et al., 1960; Watts and Strogatz, 1998; Barabási et al., 1999; Betzel et al., 2017; Váša and Mišić, 2022). The effect on network topology due to spatial constraints and the metabolic cost of establishing connections becomes increasingly important when utilizing different network models for the study of functional outcomes resulting from structural changes due to adaptive or maladaptive network responses, such as those occurring as healthy networks are subject to disease (Zhang et al., 2019; Akiyama et al., 2019; Garone et al., 2021).

To alleviate these shortcomings one can adopt spatially embedded generative network models as the basis for network generation (Waxman, 1988; Kaiser and Hilgetag, 2004). These models capture the underlying principles of neural network formation through an emphasis on the importance of spatial distributions of neurons and the wiring cost associated with axon development (Chen et al., 2006; Onesto et al., 2017). In simplified terms, wiring cost principles propose that axon development can be reduced to a set of rules governed by an optimization to reduce global wiring length while maintaining a high level of computational efficiency combined with spatial constraints on component placement (Cherniak, 1994; Ahn et al., 2006; Chen et al., 2006; Hayward et al., 2023).

Wiring cost optimization has been suggested to aim at reducing the metabolic consumption during axon formation and maintenance over time, which is especially true for long-range connections which are considered costly to both establish and maintain. To do so, neurons self-organize into clusters of neurons with similar roles leading to highly interconnected modules of segregated function, balanced by a few long-range connections to increase computational efficiency (Cherniak, 1994; Hayward et al., 2023). This results in a cost-benefit trade-off on a macro- and microscale (Kaiser and Hilgetag, 2006; Goulas et al., 2019). Long-range connections, which may seem energetically unfavorable, have the role of facilitating a reduction in average path lengths, thereby increasing the speed of information transfer across the networks (Kaiser and Hilgetag, 2006). Previous work has shown that spatial network models can capture the structures of neural networks both in vivo (Kaiser and Hilgetag, 2004) and in vitro (Onesto et al., 2017). The use of these models therefore allows for the study of a variety of mechanisms in a computationally efficient manner, such as the effect of spatial clustering on information transfer (Onesto et al., 2017; Gentile, 2024) or changes to network structures based on different wiring probability functions (Kaiser and Hilgetag, 2004; Gentile, 2021; Akarca et al., 2025).

Here we adopted spatially embedded generative network models to investigate the effects of simplified wiring cost principles combined with spatial constraints on network formation. We included two connection probability functions, one favoring short to intermediate connectivity (exponential), and one favoring short-range connectivity with a heavy tail (log-normal). Two aspects of interest were investigated, mainly how changes in spatial distributions alter the resulting networks topology in combination with different wiring probabilities, and how pruning of long-range connections changes network function. Specifically, we investigated how changes to spatial conformations, corresponding to changes to cell adhesion leading to different sizes of spatial clusters, would affect the resulting network topology, in combination with different wiring probabilities. Our choice of wiring probabilities were made to capture the protective effect long-range connections may have on network function in comparison to wiring probabilities lacking long-range connections inspired by the reduction of axon outgrowth and increased branching found in neurodegenerative disease. By doing so, we intended to determine how these differences influence the resulting network topology, thereby improving our understanding of how compensatory changes in regards to the formation and maintenance of connections in neural network topologies are altered following perturbations to network. This offers novel insight into the mechanisms underpinning neurodegenerative diseases such as ALS, and how these may ultimately lead to a loss of information transfer and network function.

## Results

In the models used in this study neurons were modeled as points, termed nodes, embedded in a Euclidean space. The connections between these nodes were meant to capture the connections between neurons (e.g. physical connections such as axons or correlations in activity between regions or sub-units). We first initialized node placements before connecting each node using one of two wiring probability functions.

### Node placement

Crucial to facilitating optimal neural activity, the neurons’ physical placement in relation to each other determines their probability of making a connection, tuning the onset and presence of sustained dynamical activity. To ascertain how increased spatial cluster sizes (decreasing number of initially placed nodes) alter network hallmarks, we introduced spatial constraints to node placement leading to clusters of nodes of varying size for each spatial conformation.

We initialized the systems by generating 100 different placements for each spatial conformation. One thousand nodes were placed in the unit square for each system, with all or a subset of nodes placed uniformly distributed in the available area. For each spatial conformation, *N*_init_ equal to either 1000, 200, 100, 50 or 20 nodes were initially placed. Any remaining nodes were uniformly assigned to an initially placed node, and the assigned initially placed node was used to represent a new frame of origin. To determine the distance from the chosen frame of origin, a power-law was used with *α* = 3*/*2. The radial angles were chosen at random following a uniform probability. This resulted in systems with different spatial constraints and varying node densities around the initially placed nodes *N*_init_ as shown in Fig. 1. The size of the spatial clusters were expected to follow 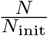 resulting in a mean size of 5 (200), 10 (100), 20 (50), 50 (20) or 1000 (1) nodes (*N*_init_), corresponding to different regimes of clustering found in in vitro studies of neural networks (Okujeni and Egert, 2019b). Due to the distance from the initially placed node being determined by a power-law, most nodes were expected to be placed close to the initial node with a non-zero probability of a node being found at a larger distance (**Fig. 1A–E**). The resulting systems showed an increasing amount of unoccupied area as the distribution of nodes shifted toward fewer and larger spatial clusters (**Fig. 1F–J**).

**Fig. 1.**
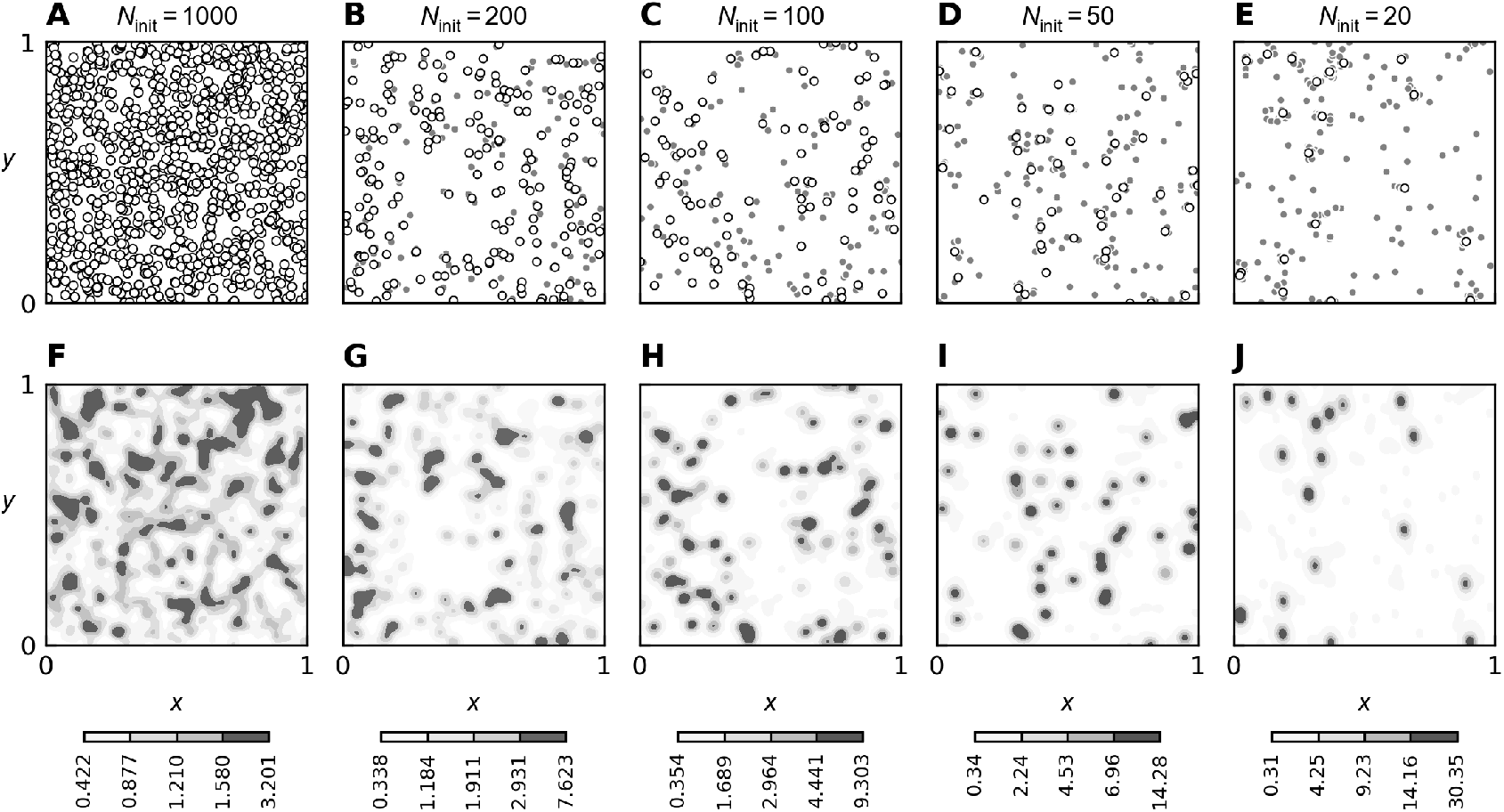
Example of placements of nodes in the unit square for each spatial distribution with corresponding kernel density estimate found using a Gaussian kernel. **(A–E)** The node distribution in the unit square for decreasing number of initially placed nodes (*N*_init_ = 1000, 200, 100, 50 and 20, respectfully). **(F–J)** The corresponding kernel density estimate for each node distribution. A total of *N* = 1000 nodes were placed in the unit square embedded on a torus for all distributions. Initially placed nodes (white circles with black borders) were placed uniformly in the available space, whereas remaining nodes (gray circles with white borders) were distributed around the initially placed nodes. The expected cluster sizes were given by 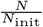 . In the density estimates a darker hue of gray indicates stronger spatial clustering of nodes. Note the difference in scale for the density distributions.

### Spatial Clustering introduces nodes with strong local connectivity

To further determine how the spatial distribution changed local interactions for increased spatial clustering, we estimated the probability density *P* (*d*) and cumulative distribution function CDF of finding a connection as a function of the Euclidean distance to the nearest 50 neighbors (**Fig. 2A–B**). All systems with spatial clustering showed a strong affinity for the nearest node to be found in close proximity in contrast to the uniformly distributed systems. At a distance *d* ≪ 0.01, the distributions were strongly influenced by the expected cluster size, and in all systems with clustering there was a peak close to the smallest allowed distances from the initial node. This peak was more prominent for larger cluster sizes. For the systems with *N*_init_ equal to 200, 100 or 50 a bimodal distribution was visible (**Fig. 2A**). This suggests that systems consisted of a densely packed core with interactions to distant nodes corresponding more closely to a uniform distribution. Here, the expected cluster sizes were lower than the number of neighbors of interest and distant clusters contributed to these similarly to randomly placed nodes. This bimodal distribution was visible for all systems outside of the uniformly distributed system with the exception of the system with *N*_init_ = 20, likely due to the number of nearest neighbors considered corresponding with the expected cluster size.

**Fig. 2.**
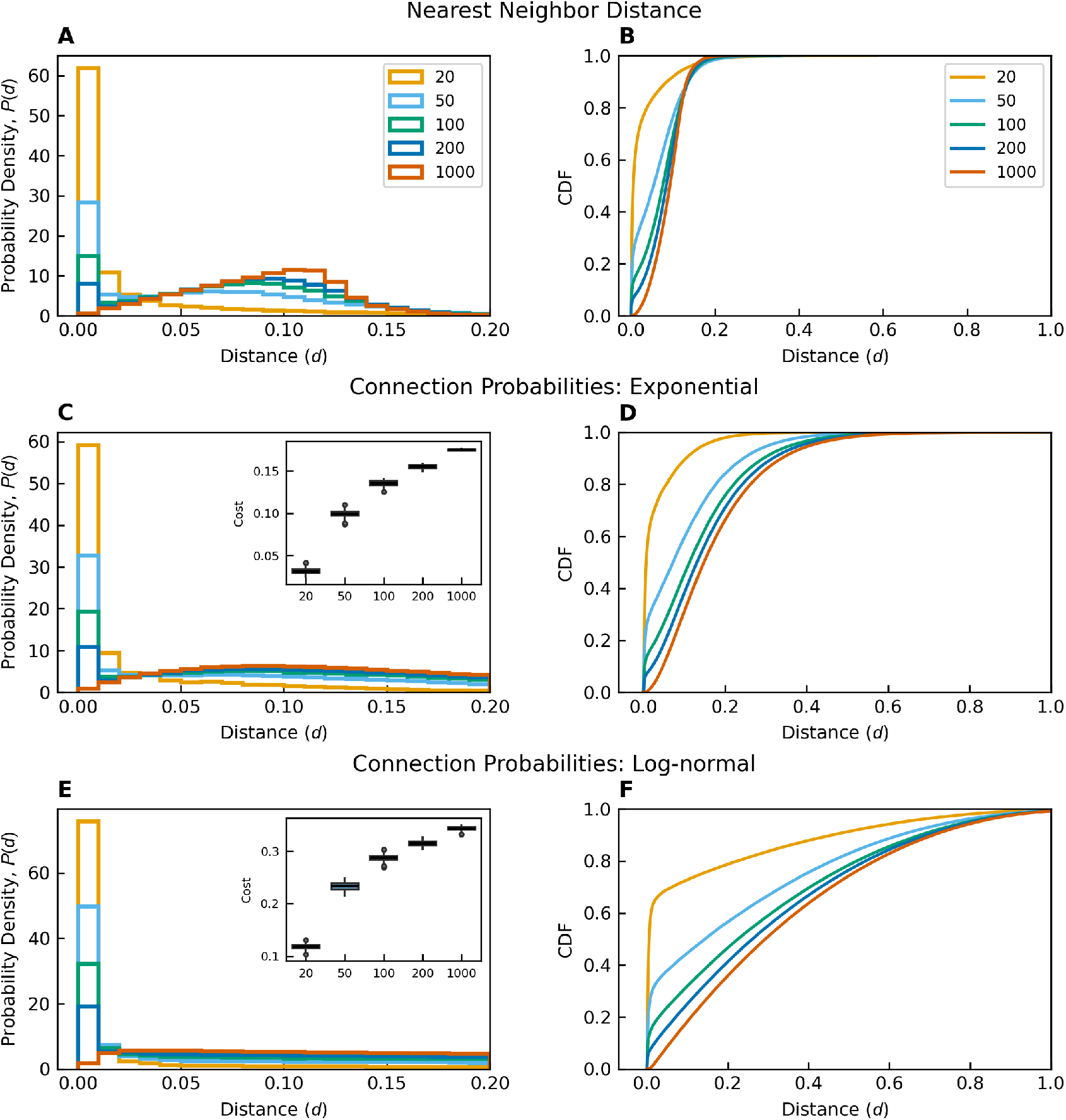
Pooled nearest neighbor probabilities and connection probabilities for the exponential and log normal wiring probability functions with corresponding cumulative distribution functions. **(A)** Probability densities (*P* (*d*)) of finding the 50 nearest neighbors at a given Euclidean distance *d* for all spatial conformations at short distances (*d* ≤0.2) with **(B)** the corresponding cumulative distribution function (CDF) at distances (*d* ≤1.0). Expected cluster sizes decreases with increasing spatial clustering, such that for *N*_init_ = 20 initially placed node the expected number of nodes distributed around each initially placed node was approximately 50, while one for *N*_init_ = 1000. **(C)** The *P* (*d*) of finding a connection between two nodes at a distance *d* for the exponential wiring probability function and **(D)** corresponding CDF. Similarly, **(E)** the *P* (*d*) and **(F)** CDF for the probability of finding a connection between two nodes for the log-normal wiring probability function. Insets in **(C)** and **(E)** shows the wiring cost for the respective wiring probability. Networks generated using either wiring probability function had a large peak at very short *d*, except *N*_init_ = 1000. For *N*_init_ = 20 most connections were within *d* ≤ 0.2 for the exponential wiring probability whereas the log normal wiring probability showed less of a dependence on local connections outside of the immediate cluster. The wiring cost increased with reduced spatial clustering, and was larger overall for the log-normal wiring probability as compared to the exponential. All systems had *N* = 1000 nodes, and 100 instances were generated for each spatial conformation and wiring probability. Legends are shared between figures.

For each spatial conformation, we generated networks using tuning parameters for both the exponential and log-normal wiring probabilities chosen such that the mean degree ⟨*k*⟩ was ⟨*k*⟩ ≈50. By calculating the probability *P* (*d*) of finding a connection at distance *d*, we found that both systems had a high affinity for short-range connections when spatial constraints (i.e. clustering) were introduced (**Fig. 2C– E**). Similarly to the nearest neighbor distances, a bimodal peak was visible for the exponential wiring probability. For the log-normal wiring probability, the bimodal distribution was not present to the same degree, suggesting that these networks wire either locally within clusters or irrespective of distance due to long-range connectivity as expected. This is supported by the CDF for each system where the exponential networks were shown to have most connections within *d <* 0.4 while the log-normal networks had a knee at low distances and a slower increase from 0 to *d <* 1.0 (**Fig. 2D–F**).

In both systems, the *P* (*d*) had a probability distribution at short ranges determined by an increased probability of interactions for lower *N*_init_ due to the increased number of neighbors within spatial clusters. By introducing spatial constraints, the distance between nodes was reduced leading to an overall reduction in wiring cost as seen in the insets in **Fig. 2C, E**. As expected, due to the heavy-tailed nature of the log-normal probability distribution, the wiring cost for these networks were found to be higher compared to the exponential wiring probability for all spatial constraint .

### Spatial clustering promotes modularity and small-worldness

Through the introduction of clustering, the variation in average number of connections increased with decreasing *N*_init_ (**Fig. 3A**). By reducing the inter-nodal distances by means of clustering, more connections were able to form between nodes due to the increased connection probability following the decrease in distance to an increasing number of neighbors within clusters. This in turn made each system more sensitive to local differences in spatial conformations (**Fig. 2**). These changes were more prominent for the exponential wiring probability, suggesting that this wiring probability was more sensitive to neighboring clusters whereas the log-normal systems maintained connectivity primarily within the cluster. Both diameter and average shortest distance ⟨*d*⟩ were larger in systems with large spatial cluster sizes (**Fig. 3B–C**). The log-normal systems had a lower diameter and ⟨*d*⟩ compared to the exponential systems for all spatial distributions, as expected due to the presence of long-range connections.

**Fig. 3.**
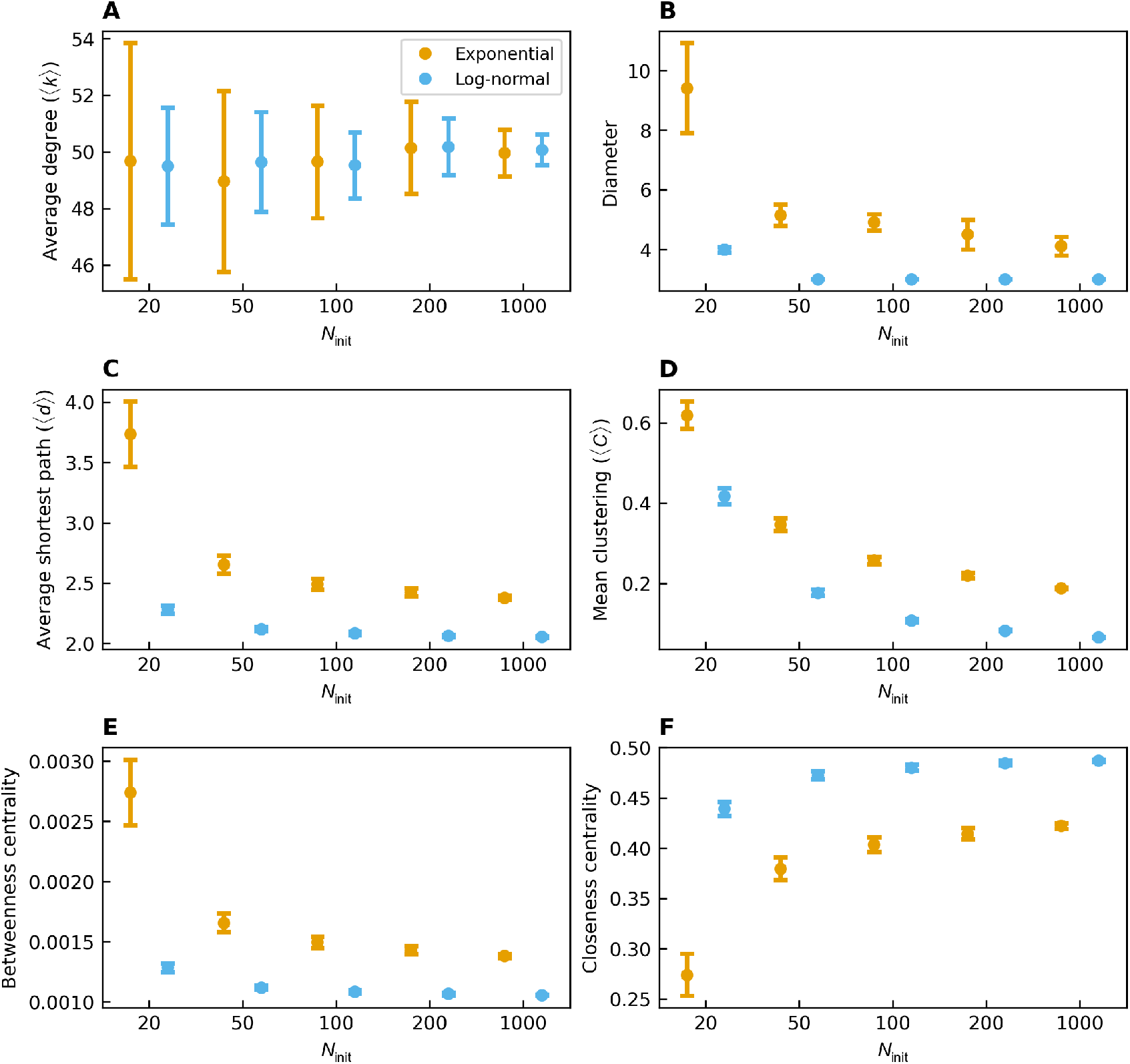
Parameters describing common network characteristics with networks tuned to have a mean degree of ⟨*k*⟩ = 50 for increasing number of initially placed nodes *N*_init_. **(A)** Average degree ⟨*k*⟩, **(B)** network diameter, **(C)** average shortest path ⟨*d*⟩, **(D)** mean clustering ⟨*C*⟩, **(E)** the betweenness centrality, and **(E)** closeness centrality for each spatial conformation. For all values, except closeness centrality, the value decreased with reduced spatial clustering. The exponential wiring probability (orange) showed higher values as compared to the log-normal wiring probability (blue), for all systems with the same spatial configurations, outside of the closeness centrality. The exponential systems had a stronger dependency on the spatial distribution of nodes, especially for *N*_init_ = 20. All systems had *N* = 1000 nodes, and 100 instances were generated for each spatial conformation and wiring probability. Each figure shows the mean generated from all realizations of each configuration with error bars showing the standard deviation. Legends are shared between figures.

In both systems, increasing spatial clustering resulted in an increase in average clustering ⟨*C*⟩ for the resulting networks (**Fig. 3D**). However, for the exponential systems the introduction of spatial constraints led to an overall larger increase in ⟨*C*⟩ compared to the log-normal systems. The introduction of spatial clustering led to an increase in local connectivity, resulting in increased ⟨*C*⟩ in both systems, with a decrease in ⟨*d*⟩ due to reduced path lengths within spatial clusters. By introducing long-range connections, the probability of a node at either end of a long-range connection to have a neighbor with a connection to the same node was reduced, suggesting that the increase in clustering was driven by short-range connectivity within spatial clusters following the reduced *N*_init_.

In the case of the exponential wiring probability, the betweennes centrality decreased and closeness centrality increased with increasing *N*_init_ (**Fig. 3E–F**). The same trend was visible for the log-normal connection probability, but with changes being less prominent. Overall, the closeness centrality was higher for all log-normal systems, whereas the betweennes centrality was higher for the exponential systems. This suggests that the systems connected by the exponential wiring cost were more dependent on a few nodes for traversing the network while most nodes were distant from each other. Especially for *N*_init_ = 20, where the betweenness centrality and closeness centrality displayed a strong reliance on a few nodes for communication and a larger distance between all nodes in the networks. Conversely, the networks created using the log-normal wiring probability were less dependent on central nodes and most nodes in the network were fairly close due to the presence of a larger number of long-range connections, even with varying *N*_init_. This was supported by the lower average path length and diameter, which did not change as much for the log-normal wiring probability compared to the exponential, even for the most clustered spatial conformations.

For low spatial clustering densities, neither wiring probability were found to produce small-world networks according to the small-world propensity *ϕ* (**Fig. 4A**). Only in systems containing large clusters (*N*_init_ ≤ 50 for the exponential systems and *N*_init_ = 20 for the log-normal systems) did we find networks which were considered to be small-world, indicating that spatial clustering in the conformation of nodes in the networks was necessary to reach a sufficient balance between clustering and short average path lengths to be termed small-world. The log-normal wiring connection probabilities have a steep initial decline with a sustained, albeit low, probability of making long-range connections, whereas the exponential systems favor short to intermediate connections. The probability for a connection at a given distance can be seen in Appendix A Fig. A.1. This affinity for local interactions resulted in a sufficiently large ⟨*C*⟩ leading to small-worldness for networks created using the exponential wiring probability for *N*_init_ ≤ 50 (**Fig. 3D**). In opposition, for the log-normal wiring probability, the decreased ⟨*d*⟩ due to long-range connections was not sufficient to facilitate an increase in *ϕ*, as the presence of these connections had a detrimental effect on ⟨*C*⟩ (**Fig. 3C**). The same was seen for systems with expected spatial clusters much smaller than ⟨*k*⟩ . Here connections were forced beyond the local vicinity of the respective spatial cluster, resulting in an overall reduction of ⟨*C*⟩ due to the decreased probability of two neighbors sharing a connection at longer distances from the immediate node. This was further supported by the degree assortativity, which was higher for all exponential wiring probabilities compared to the log-normal. Furthermore, we found that the degree assortativity increased with decreasing *N*_init_, as expected as an increasing number of connections were recruited within the spatial clusters making it more likely that all nodes within a cluster were connected and therefore share the same number of connections.

**Fig. 4.**
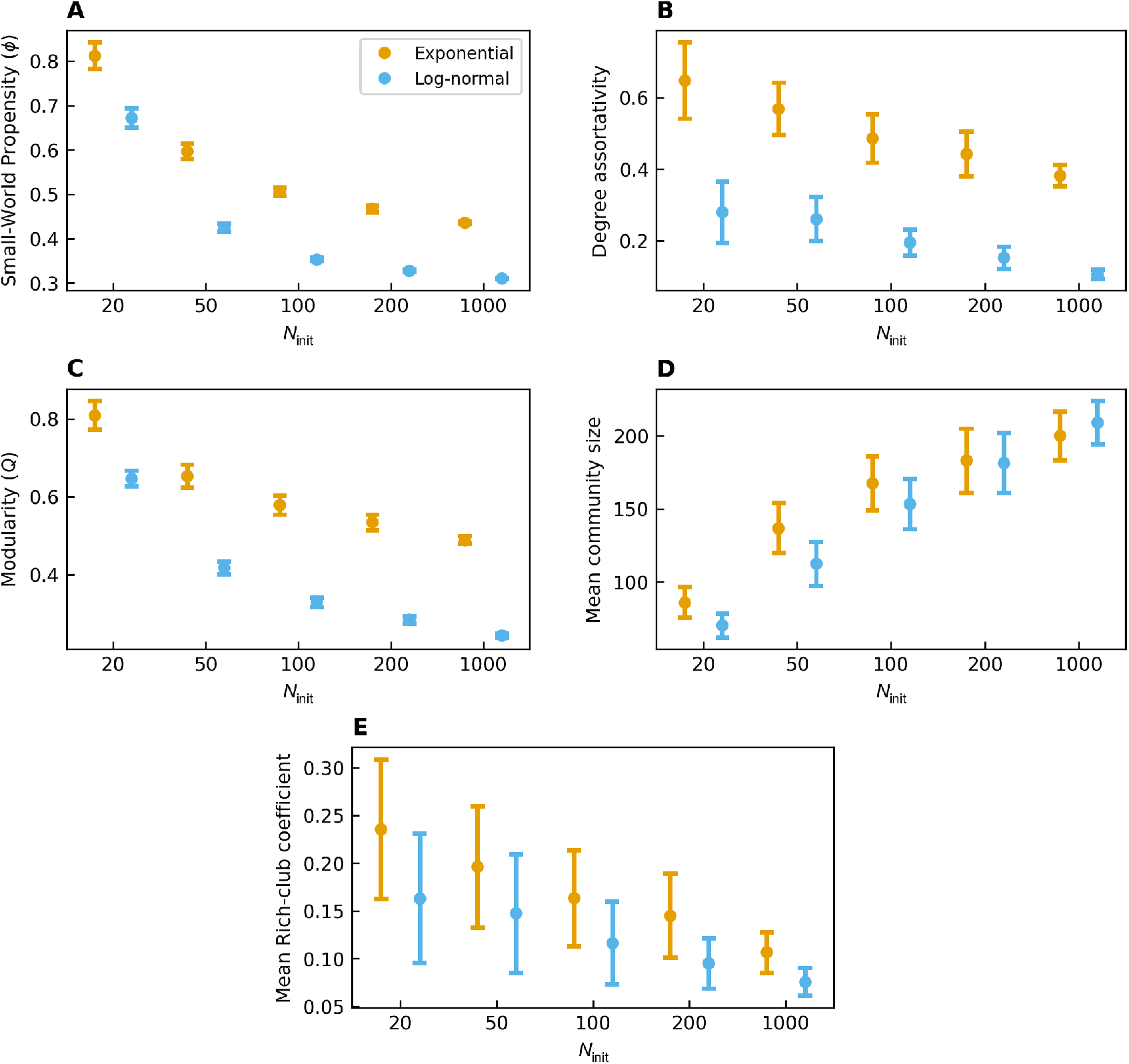
Parameters describing network characteristics related to network organization for networks tuned to have a mean degree of ⟨*k*⟩ = 50 and increasing number of initially placed nodes *N*_init_. **(A)** Small-world propensity *ϕ*, **(B)** degree assortativity, **(C)** *Q* modularity and **(D)** corresponding mean community size, and **(E)** mean rich-club coefficient. All values increase with decreasing *N*_init_ (increasing cluster size) except community size which decreases. Both wiring probabilities showed small-worldness (*ϕ >* 0.6) only for the most clustered systems; *N*_init_ ≤ 50 for exponential wiring probability (orange) and *N*_init_ ≤ 20 for the log-normal wiring probability (blue). Similarly, modularity was strongest in the systems with the strongest spatial constraints. Networks generated using the log-normal wiring probability had fewer intermediate connections compared to the exponential wiring probability leading to a reduction in connections between adjacent spatial clusters, with the presence of long-range connections reducing clustering due to a reduced probability for nodes at either end of a long-range connection to share neighbors. This leads to a disruption in community structures and small-worldness, even at low *N*_init_. All systems had *N* = 1000 nodes, and 100 instances were generated for each spatial conformation and wiring probability. Each figure shows the mean generated from all realizations of each configuration with error bars showing the standard deviation. Legends are shared between figures.

To ascertain how increased spatial clustering would affect separation into communities, we determined network partitions for all networks using *Q* modularity (Blondel et al., 2008; Newman and Girvan, 2004). For all spatial distributions, the networks generated using the exponential wiring probability had a larger *Q* modularity compared to the log-normal, where both displayed decreasing modularity scores and increasing cluster sizes with increasing *N*_init_ (**Fig. 4C**). Both wiring probabilities showed a similar change in the number of communities with increasing *N*_init_ (**Fig. 4D**). Even so, we found that the mean community size were slightly larger for *N*_init_ = 1000 for log-normal systems, whereas they were lower from *N*_init_ ≤ 100, suggesting that the local connectivity was more contained to the immediate region of any given node for the log-normal compared to the exponential wiring probability. Similarly, for all spatial distributions, *Q* was lower for the log-normal wiring probability. This suggests that although the reduction to intermediate connections leads to a stronger separation between adjacent clusters and thereby increasing the number of detected communities, the presence of long-range connections and reduced ⟨*k*⟩ results in weaker community structures. Similarly to the degree assortativity, we found the mean rich-club coefficient increased with decreasing *N*_init_ (**Fig. 4E**). The affinity for connections to be made locally for the exponential wiring probability was further supported by the larger mean rich-club coefficient overall. In the networks generated using the log-normal wiring probability, we found that the presence of long-range connections, at the cost of intermediate connections, resulted in a decrease in most values associated with efficient computation.

### Long-range connections improves information transfer

To determine how the choice of wiring probability in combination with differences in spatial conformations change the networks′ capacity for information transfer and communication, we determined the normalized global, local and diffusion efficiencies, and communicability (**Fig. 5**). The raw and relative data can be found in Appendix B Fig. B.1. Overall we found that the networks created using the log-normal wiring probability had a higher capacity for information transfer globally, whereas networks created using the exponential wiring probability showed a stronger affinity for local information transfer. For the log-normal wiring probability, decreasing *N*_init_ reduced the capacity for global communication, but less so than for the exponential. This suggests that while spatial clustering is necessary for increasing local connectivity, by strengthening local information transfer, the presence of long-range connections is crucial to maintain efficiency in communication network wide.

**Fig. 5.**
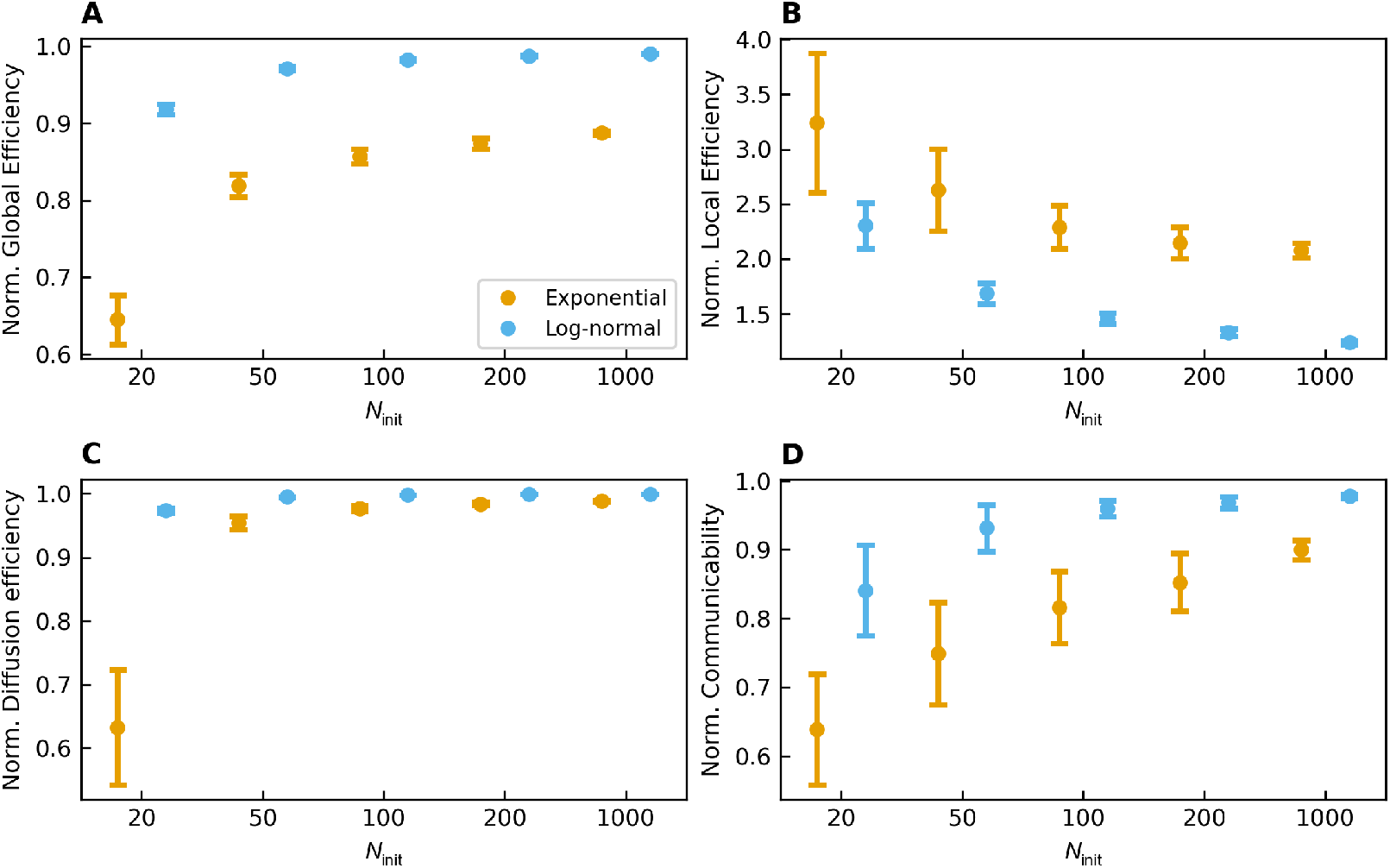
Normalized efficiency measures for all networks tuned to have a mean degree of ⟨*k*⟩ = 50 and increasing number of initially placed nodes *N*_init_. **(A)** Normalized global efficiency, **(B)** normalized local efficiency, **(C)** normalized diffusion efficiency, and **(D)** normalized communicability. For all spatial conformations, the log-normal wiring probability (blue) show better capacity for global communication whereas he exponential wiring probability (orange) show better local communication. With decreasing *N*_init_ (increasing cluster sizes) both networks to displayed an increase in local efficiency leading to a larger capacity for local specialization. For the networks generated using the exponential wiring probability this came with a large decrease in global integration, whereas this was maintained in the networks generated using the log-normal wiring probability due to the presence of long-range connections. All systems had *N* = 1000 nodes, and 100 instances were generated for each spatial conformation and wiring probability. Each figure shows the mean generated from all realizations of each configuration with error bars showing the standard deviation. Legends are shared between figures.

### A. Pruning leads to a relative increased local connectivity

Different network structures have different resilience towards perturbations to the network. By pruning the networks we aimed to elucidate how targeted pruning changes network structure, and subsequently function. Pruning was performed by removing the longest connections until the desired pruning fraction was reached for networks generated with *N*_init_ = 20. The choice of pruning long-range connections was made to simulate the gradual breakdown of costly connections in the networks. Only fully connected networks were analyzed and used for the end results to avoid issues related to unconnected networks. This resulted in a slight decrease in the total number for networks generated using the log-normal wiring probability, with 92 fully connected networks at a pruning degree of 0.05 and 79 remaining networks at a pruning degree of 0.1. The networks generated using the exponential wiring probability were excluded from this analysis as the number of connected networks were insufficient to be considered. The number of connected and unconnected networks for each wiring probability can be seen in Appendix B Table 2. For increasing pruning fraction we found that most calculated metrics increased, including the network diameter, ⟨*d*⟩, ⟨*C*⟩, *ϕ*, betweenness centrality, degree assortativity, *Q*, community size and rich-club coefficient, while density, ⟨*k*⟩ and the closeness centrality decrease (**Fig. 6**). By removing the longest connections the networks became increasingly reliant on the remaining long-range connections, while all in nodes became more distant as shown by the increase in betweenness centrality and decrease in closeness centrality. As such, the networks became more strongly segregated into modules with a relative increase in ⟨*C*⟩ . This increase in ⟨*C*⟩ was a result of pruning nodes which were more likely to not have neighboring nodes which were connected to itself. Furthermore, the pruning forced the networks to increasingly rely on wiring within spatial clusters. Even so, we found little change in the rich-club coefficient, suggesting that hubs in our networks were not driven by long-range connections, but reliant upon connectivity within spatial clusters.

We determined how the networkś capacity for information transfer and communication changed due to pruning by determining the normalized global, local and diffusion efficiencies, and communicability for increasing pruning (**Fig. 7A–D**). We found that measures related to global information transfer were decreased, whereas local efficiency was increased. While we did not change or add connections, the relative wiring both locally and globally changed with targeted pruning. Due to the targeting of long-range connections, the resulting networks had a decrease in capacity for global information transfer corresponding with increased path lengths and diameter of the networks, whereas local connections were strengthened as seen in both increased clustering and local efficiency.

**Fig. 6.**
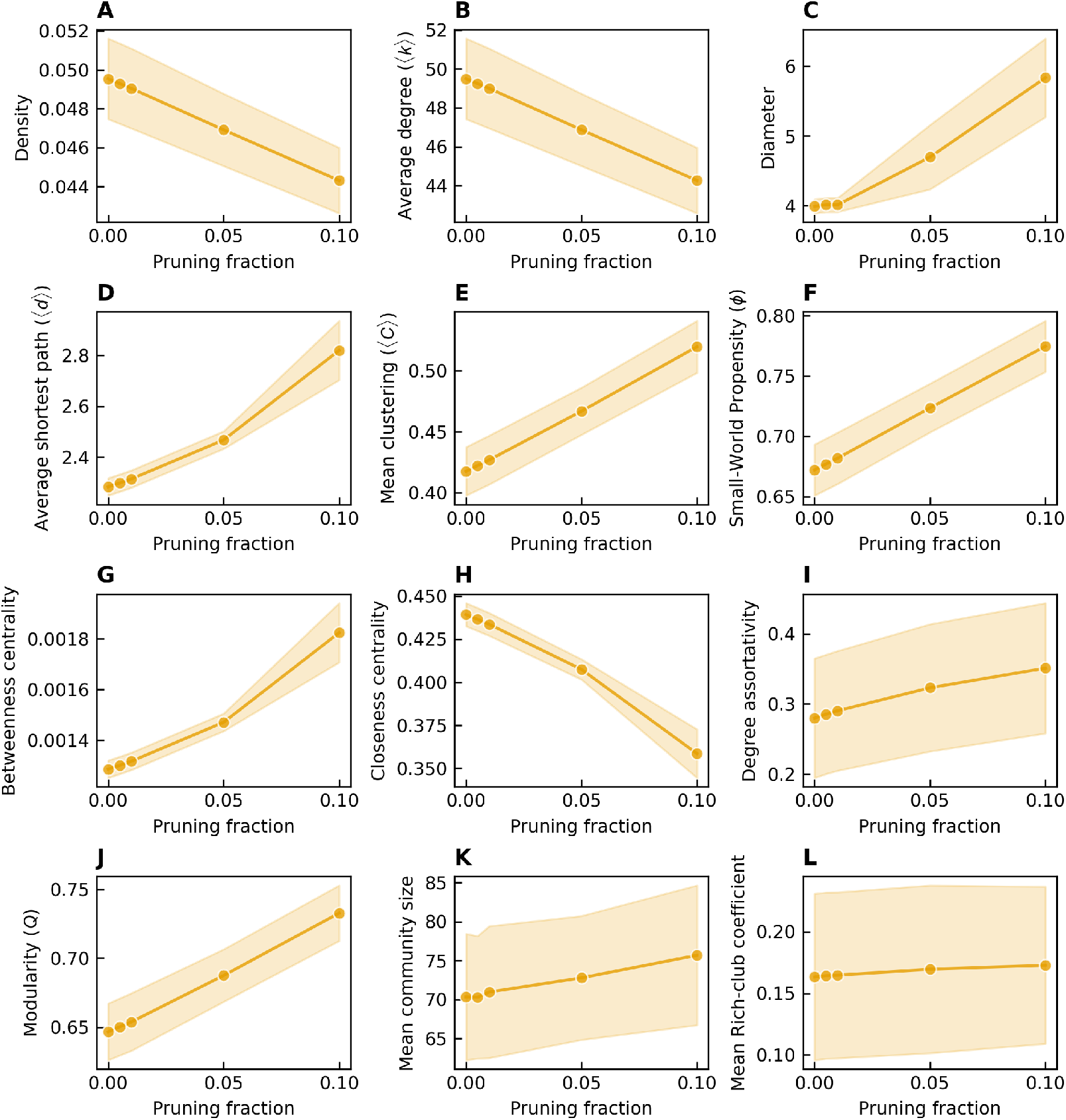
Parameters describing network characteristics for networks generated using the log-normal wiring probability and initially placed nodes *N*_init_ = 20 at varying pruning degrees. **(A)** Network density, **(B)** average degree ⟨*k*⟩, **(C)** diameter, **(D)** average shortest path ⟨*d*⟩, **(E)** mean clustering ⟨*C*⟩, **(F)** small-world propensity *ϕ*, **(G)** betweenness centrality, **(H)** closeness centrality, **(I)** degree assortativity, **(J)** *Q* modularity and **(K)** corresponding mean community size, and **(L)** mean rich-club coefficient for increasing pruning fraction. All systems have *N* = 1000 nodes and the same networks were pruned at varying degree. Only fully connected networks were analyzed leading to a slight reduction in the number of networks for the strongest pruning degrees (92 networks at a pruning degree of 0.05 and 79 networks at a pruning degree of 0.1.). Each figure shows the mean generated from all networks for each pruning degree with the shaded area showing the standard deviation.

**Fig. 7.**
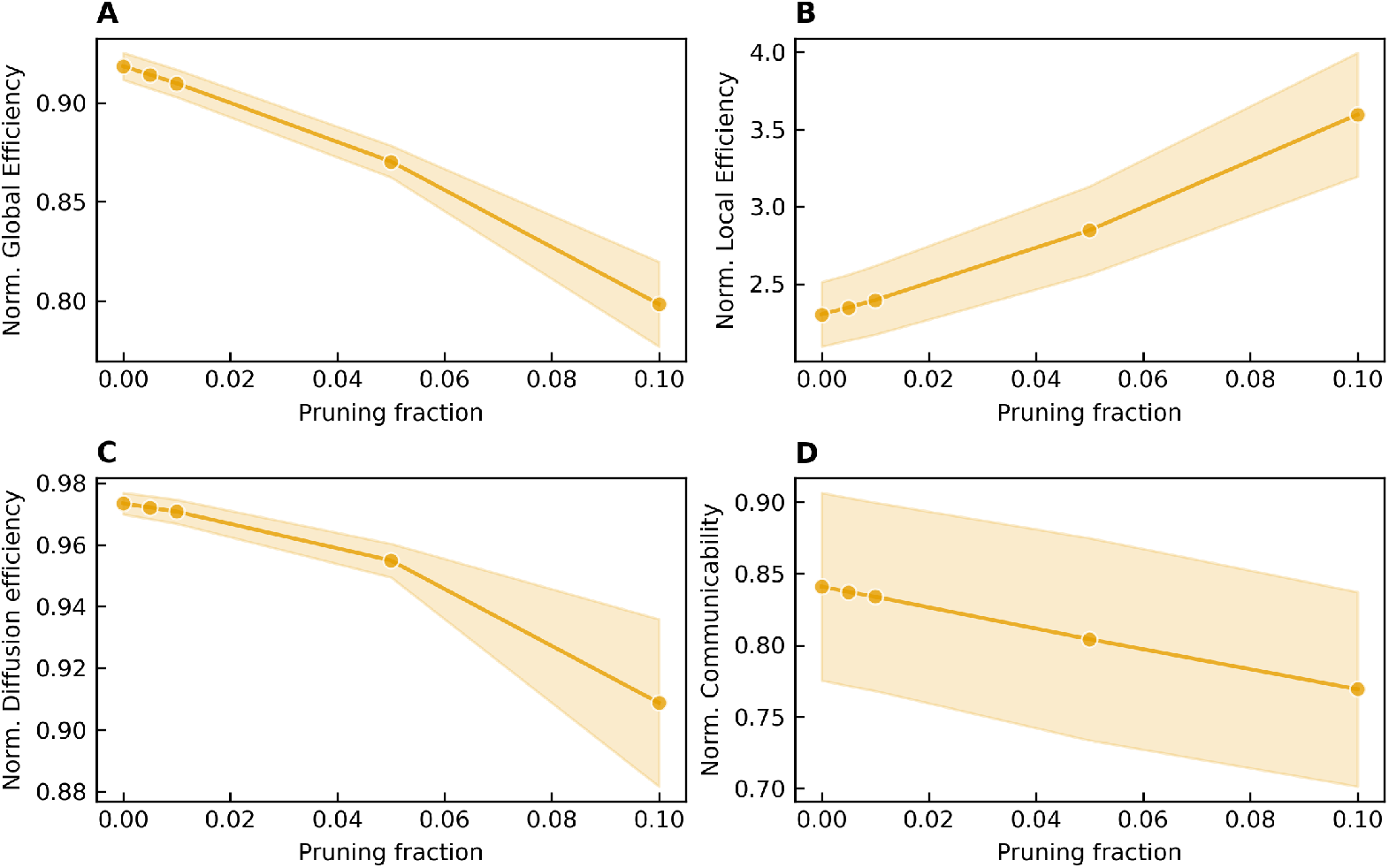
Normalized efficiency measures for networks generated using the log-normal wiring probability and initial nodes *N*_init_ = 20 at varying pruning degrees of long range connections. **(A)** Normalized global efficiency, **(B)** normalized local efficiency, **(C)** normalized diffusion efficiency, and **(D)** normalized communicability. Metrics related to global efficiency and information transfer are reduced with increasing pruning of long-range connections whereas local efficiency is strengthened. With increased pruning measures related to global efficiency and information transfer are reduced while local efficiency is increased. All systems have *N* = 1000 nodes and the same networks were pruned at varying degree. Only fully connected networks were analyzed leading to a slight reduction in the number of networks for the strongest pruning degrees (92 networks at a pruning degree of 0.05 and 79 networks at a pruning degree of 0.1.). Each figure shows the mean generated from all networks for each pruning degree with the shaded area showing the standard deviation.

## Discussion

In this study, we investigated how the choice of wiring probability function alters structural outcomes for networks with a variety of spatial conformations when generating networks using a simple spatial network approach inspired by wiring cost principles, and how these wiring principles affect the networks ability to maintain efficient communication in the face of perturbations. We found that adding spatial constraints in the form of clustering to the node placement of the spatial models improves network properties associated with improved functional outcomes in biological neural networks. The introduction of larger spatial clusters of nodes leads to a reduction in the overall wiring cost while also increasing network clustering, resulting in networks considered to be modular and small-world, combined with increased local efficiency of information transfer. Even with strong spatial clustering, we found that global communication was retained for networks with long-range connections. Our findings further corroborate the hypothesis, that the small-worldness of these systems depends on the majority of connections being constrained within the respective cluster to which the node was initially placed, with a high probability of neighbors sharing connections. This necessitates that the networks are sufficiently sparse to limit connections to within the local vicinity of the node, yet share a limited number of long-range connections to distant modules in the network to balance local integration with global computational efficiency.

Networks generated using either connection probability were only considered small-world when the nodes of the network were constrained to sufficiently large spatial clustering. In accordance with previous work, we found that networks generated using a heavy-tailed connection probability, such as the log-normal connection probability, struggled to achieve small-worldness without sufficiently large constraints in the spatial domain for sparse networks compared to the exponential distribution (Kaiser and Hilgetag, 2004). The presence of long-range connections in the networks resulted in an overall reduction in average path length, but also impacted network clustering. The possibility of adding connections to just about any node in the system reduces the probability of a node connected distally to be connected to a second neighboring node, as the nodes they connect to have a lower probability of making a connection to a neighbor in the local vicinity of the initial node, thereby reducing the network clustering. As such, the long-range connections had a negative effect on both small-worldness and modularity. The networks generated using the exponential wiring probability were more readily pushed into a small-world state, but in contrast to previous work, we found that sufficient spatial clustering was necessary also for this wiring probability to be considered small-world (Kaiser and Hilgetag, 2004; Onesto et al., 2017, 2019; Gentile, 2021). Overall the networks generated using the log-normal wiring probability had lower modularity scores as compared to the exponential, but also separate into a slightly larger number of communities for spatially clustered conformations. This suggest that the reduction in intermediate ranged connections resulted in segregation into fewer distinct communities. Even so, the overall modularity score was lower, suggesting that standard null models may not be able to capture the intricacies of spatially embedded models, and more representative modularity scores could be determined using null models better suited for the task (Váša and Mišić, 2022).

The dynamical range and onset of activity in neuronal networks is inherently dependent on the structure of the network on which the activity propagates. This dependence has been seen both in computational studies and in in vitro models of neural networks (Massobrio et al., 2015; Yamamoto et al., 2018; Arvin et al., 2022). Through constraints on neuronal organization in the spatial domain, combined with wiring optimization, these networks have been shown to develop a high local clustering with long-range connections, resulting in a reduced number of steps necessary to traverse the network (Cherniak, 1994; Kaiser and Hilgetag, 2006; Hayward et al., 2023). The high degree of local clustering aids in increasing local feedback loops, whereas the network wide interconnectedness arising from long-range connections facilitate global interactions and synchrony in the networks, necessary for processing information (Markov et al., 2013; Okujeni et al., 2017; Arvin et al., 2022; Avramiea et al., 2022; Antonello et al., 2022).

Through using spatial network models, we found that structures associated with these dynamical ranges appeared only as a consequence of high degrees of spatial clustering of nodes in the network, irrespective of the chosen wiring probability. With increasing spatial clustering and decreasing internodal distances, clustering and modularity increased. This indicates a shift towards a globally more segregated community structure with locally integrated nodes, in agreement with findings stating that component placement plays an integral role in reducing the overall path length through local integration while also supporting formation of structures associated with diversified and sustained activity (Cherniak, 1994; Okujeni et al., 2017). These findings were further supported by the increase in efficiency of information transfer locally for systems with larger degrees of spatial clustering, whereas the global efficiency was reduced. Notably, the effect on the reduction in global information transfer for increasing spatial clustering was found to be more pronounced for the networks lacking long-range connections. Networks with long-range connections were less affected by changes to spatial distributions of nodes, highlighting the importance of these connections for maintaining function and network wide communication.

We found that even a slight change in local connectivity with the introduction of long-range connections disrupts features considered important for computation in biological neural networks. Although such mechanisms may seem trivial, the apparent contradiction that the presence of long-range connections reduces small-worldness, modularity and local integration helps shed light onto mechanisms underlying disease or aging. In progressive neurodegenerative diseases such as ALS, the networks in vitro have been shown to undergo alterations including increased neurite branching and reduced outgrowth (Akiyama et al., 2019; Garone et al., 2021; Kollstrøm et al., 2025a). Analogous to our systems, an increase in neurite branching and local clustering combined with reduced long-range connections have been suggested to increase the overall small-worldness and modularity measures, all while resulting in reduced global efficiency (Rubinov and Sporns, 2010; Zhang et al., 2019). As we have previously shown in networks subject to these maladaptive conditions, such alterations can occur at different stages, ultimately affecting the activity of the networks (Kollstrøm et al., 2025a; Valderhaug et al., 2024; Weir et al., 2023, 2024; Kollstrøm et al., 2025b). In such diseases, where cell adhesion and neurite outgrowth are altered, the networks may be left vulnerable to perturbations, especially to targeted pruning of the metabolically costly long-range connections.

By studying the targeted deletion of long-range connections we gain an understanding of how these changes may alter network topologies analogous to progressive disease spread. For the networks lacking long-range connections, the majority of the networks were disconnected even at low pruning degrees. This highlights the vulnerability of such networks and the protective function long-range connections have towards perturbations, where even a low reduction in long-range connectivity leads to a break down of network structure resulting in disconnected networks, thereby removing any possibility for global communication. For the networks containing long-range connections, the networks were resilient to pruning, maintaining network wide communication, albeit with reduced efficiency. The gradual reduction of long-range connections forced the networks to rely increasingly on a select few nodes for global communication, while increasing the overall path lengths. This in turn resulted in network topologies often related to efficient communication being increased, including small-worldness and modularity. Furthermore, we found that the networks ′ ability to transfer information globally was hindered, with increasing reliance on information integration locally within spatial clusters, in correspondence with previous findings (Zhang et al., 2019). However, the increase in small-worldness, modularity and local efficiency in the current study was not the result of new connections, but rather the result of an increase in the relative number of local connections due to pruning of long-range connections leading to an increased reliance on local integration of information. As such, we found that changes often related to maladaptive responses in networks may also appear as a natural consequence of the deletion of long-range connections. Although these changes may seem favorable for the onset and maintenance of activity in the networks, they put additional strain on local circuitry and remaining long-range connections, which in turn may further disease cascades and network breakdown (Fiskum et al., 2026).

In summary, we have modeled how changes to spatial conformations change the resulting networks’ ability to efficiently communicate for networks with or lacking long-range connections. We found that spatial clustering was necessary, but not sufficient for functional efficiency, leading to an increase in local integration while reducing global communication. The presence of long-range connections was shown to reduce some topological metrics related to onset of activity and dynamical range, but preserved function even for networks with large spatial clustering, maintaining connections necessary for global synchrony and communication. Furthermore, we found that the loss of long-range connections could appear as an improvement in topology while degrading communication, allowing for novel insight into the effects of spatial clustering, wiring probability, and targeted deletion of connections. By combining these methods with an extensive list of network metrics, we can understand how disease spreads and establish targets for intervention, especially in networks subject to alterations in mechanisms related to cell adhesion, migration, and axon outgrowth, and how these mechanisms leaves networks vulnerable to progressive neurodegenerative disease or perturbations.

## Methods

### Network generation

To generate networks two steps were performed. Firstly, nodes were assigned a position in a given space to act as a basis for the spatial embedding for each network. Secondly, nodes were connected through the adoption of a statistical wiring probability function. Two facets of interest were studied, namely how network topology changes due to the introduction of spatial constraints and clustering for the spatial embedding, and how a pruning of long-range connections alter network hallmarks.

#### Node placement

Nodes were placed on a Euclidean plane in a one-by-one unit square. The nodes were placed on the plane following either a uniform distribution of the nodes in the available area, or with nodes in clusters. In the case of the clustered distributions, an initial number of nodes (200, 100, 50 or 20) were uniformly distributed in the plane with subsequent nodes assigned an initially placed node. Allocation of nodes to a given cluster (i.e., initially placed node) was performed by uniformly distributing the remaining nodes among the initially placed nodes to ensure that assignment was random while clusters were kept at a comparable size. During node placement, the plane was treated as a torus, resulting in periodic boundary conditions such that all nodes where bound to the unit square. A total of 1000 nodes were placed in each system, and a total of 100 systems were generated for each spatial conformation.

The distances *r* from the initially placed nodes were chosen from a power law distribution determined by the equation

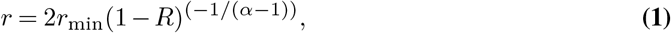

according to Newman (2005). Here *r*_min_ is the radius of the node, *R* a random number in the range [0, 1) chosen from a uniform distribution, and *α* the power law exponent. The radius *r*_min_ was set to 0.0001 to reflect neurons inability to occupy the same space (Okujeni and Egert, 2019b), while *α* was chosen to be 1.5 to balance clustering around the initially placed nodes. The radial directions of the nodes were chosen from a uniform distribution of angles in the range [0, 2*π*). In all cases, the position of each node was redrawn if the distance between any pair of nodes was less than 2*r*_min_ to avoid spatial overlap.

To ensure that streams of random numbers were not repeated, random numbers were handled according to the documentation using the Python package NumPy (Harris et al., 2020). A set of seeds were generated from an initial seed for each use case of random numbers. For each network, near independent seeds were generated to get unique streams of random numbers and avoid overlap.

#### Wiring probability functions determine connection probability by distance

Networks for each system were generated by connecting the nodes according to Waxman (1988). Whether or not an edge would occur between a given pair of nodes {*u, v*} was determined by

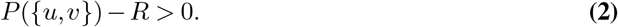

Here *R* is a random number in the range [0, 1) determined by a uniform distribution, and *P* ({ *u, v* }) the probability function determining if a connection would be made. In Eq. 2, the wiring probability function *P* ( {*u, v* }) is analogues to the wiring cost in neural networks, where the Euclidean distance *d* _{*u,v*}_ between two nodes determines the probability that a connection would occur between a set of nodes.

To determine the connection probability, two separate wiring probability functions were used for each system, an exponential and a log-normal probability function respectively. In the case of the exponential wiring probability function, *P* ({*u, v*}) was given by the equation

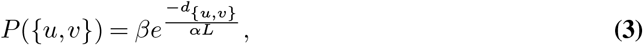

where *α, β* and *L* are tuning parameters. Although *L* can be interpreted as the largest calculated distance in the system, we kept *L* constant at 1 to avoid any differences between different spatial configurations.

In the case of the log-normal wiring probability, *P* ({*u, v*}) was given by

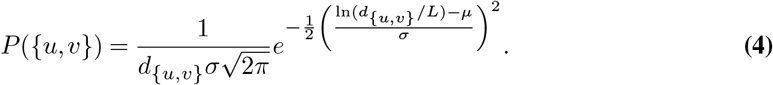

Again, *µ* and *σ* are tuning parameters used to determine the shape of the curve for the wiring probability, while *L* is defined similarly as in Eq. 3. By tuning the wiring probability functions, the number of connections established is changed. In both cases, the probability of making a connection decreases with increasing distance between nodes.

The tuning parameters were chosen such that the density was comparable for both wiring probabilities and all spatial distributions. This was done to determine the effect of spatial clustering on hallmarks of the network where the density was approximately invariant. For all simulations, *β* = 1 and *µ* = 0 were kept constant for their respective wiring probability function. The tuning parameters *α* and *σ* were determined by iterating over a range containing the chosen ⟨*k*⟩ . For each value, 5 networks were generated, and the value with the mean closest to the chosen ⟨*k*⟩ was used for the final network generations. A value of ⟨*k*⟩ ≈ 50 was chosen to correspond with the expected size of the clusters in the most dense spatial configuration. All tuning parameters can be found in Appendix B Table 1 with corresponding probability functions in Fig. A.1. During network wiring, any unconnected networks were redrawn, ensuring all networks were connected before analysis. All networks considered were binary networks. The wiring cost for each network was calculated as the mean of the Euclidean distances of all connections in the given network.

**Table 1.** Comparison of tuning parameters and mean degrees for exponential and log-normal connection probability functions. Each tuning parameter was tuned such that the mean degree was as close to 50 as possible.

| Initially placed nodes<br>($N_{\text{init}}$) | Exponential | | Log-normal | |
| --- | --- | --- | --- | --- |
|  | Tuning<br>param. | Mean<br>degree | Tuning<br>param. | Mean<br>degree |
| 20 | 0.0509 | 49.6751 | 67.2230 | 49.4933 |
| 50 | 0.0812 | 48.9675 | 34.1430 | 49.6481 |
| 100 | 0.0926 | 49.6556 | 27.8550 | 49.5276 |
| 200 | 0.0976 | 50.1345 | 25.2570 | 50.1804 |
| 1000 | 0.1024 | 49.9618 | 23.1520 | 50.0761 |

For pruning long-range connections, connections were sorted by the largest Euclidean distance between nodes and pruned (i.e. removed from the network) depending on inter-nodal distance starting with the longest distance and so on. The number of edges to remove was determined by the pruning fraction, and only networks which were still connected were used for subsequent analysis.

#### Network analysis

To classify the effects of changes to spatial distributions and choice of wiring probability function we calculated common network parameters.

#### Path length and network diameter

The average shortest path ⟨*d*⟩ is given by the average of all shortest paths, *d*_*i,j*_, between all node pairs in the network,

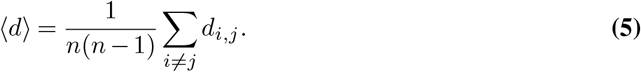

The network diameter *d*_max_ is defined as the longest shortest path or maximum eccentricity. These measures indicate the average and largest numbers of steps necessary to traverse the network from one node to the next and the largest expected distance between two nodes.

#### Clustering coefficient

To measure how well connected locally the networks were we calculated the mean clustering coefficient ⟨*C*⟩ for each network, defined as

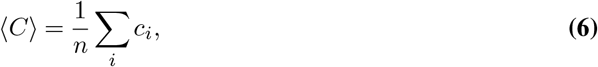

where *n* is the number of nodes in the network and *c*_*i*_ the clustering coefficient of the given node (Watts and Strogatz, 1998). The clustering of node *i* measures the number of connected neighbors indicating how many of the first degree connections of a given node are also connected connections.

#### Centrality measures

To determine how central a node is in a network we calculated the betweenness and closeness centrality (Rubinov and Sporns, 2010). The betweenness centrality measures how often a given node appears along the shortest paths of the network. The closeness centrality, defined as the inverse of the average path length, tells us how distant a node is to all other nodes of the network.

#### Small-world propensity

A small-world organization is characterized by networks having both a high degree of clustering and short average path lengths, allowing for faster information transfer and propagation across the networks (Watts and Strogatz, 1998). To measure the affinity for the networks to organize into small-world structures we estimated the small-world propensity *ϕ* for each network according to Muldoon et al. (2016). We opted for the small-world propensity as this measure has been shown to be more robust to differences between weights and binary networks and less dependent on network density compared to other measures small-worldness, making comparisons between studies easier (Telesford et al., 2011; Neal, 2017; Muldoon et al., 2016; Bassett and Bullmore, 2017). The value *ϕ* is determined by

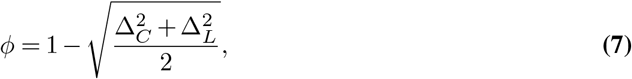

where

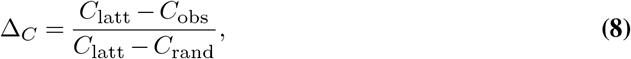

and

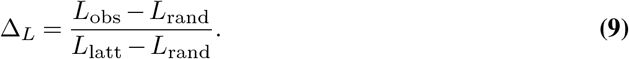

Here, *C*_obs_ and *L*_obs_ are the mean clustering coefficient and average shortest path for the observed network, and *C*_rand_, *C*_latt_, *L*_rand_ and *L*_latt_ are the mean clustering coefficient and average shortest path for equivalent (i.e. same number of nodes and edges) random (rand) and lattice (latt) networks. A total of 100 random networks were generated to determine *C*_rand_ and *L*_rand_. For *C*_latt_ and *L*_latt_ the results for a single equivalent lattice network was used due to the deterministic nature of the network. These values were used as references to determine Δ_*C*_ and Δ_*L*_. The variables Δ_*C*_ and Δ_*L*_ are limited to the range [0, 1], such that *ϕ* is constrained to the range [0, 1]. A *ϕ* of at least 0.6 is considered small-world (Muldoon et al., 2016).

#### Modular and hierarchical structures

To measure whether the networks segregated into subdivisions of densely connected modules (or communities), we calculated two separate modularity scores for each network. We first separated the network into modules using the Louvain method by varying the resolution parameter *γ* (Blondel et al., 2008). The resulting partitions were subsequently evaluated using the *Q* score for evaluating the modularity of the partitions (Newman and Girvan, 2004). For each respective modularity score we choose a range to iterate *γ* over in increments of 0.01. The modularity score was calculated 5 times for each increment to determine the *γ* value corresponding to the highest score for each respective metric in the chosen range. The final partitions were averaged over 50 repetitions for each network. For *Q*-modularity, *γ* was varied over the range[0.5, 2]. When evaluating the final *Q*-modularity a *γ*=1.0 was used.

### Degree assortativity and rich-club coefficient

To measure the affinity to which nodes of a certain degree connect to node other nodes of similar degree, we calculated the degree assortativity. The rich-club coefficient was used to measure whether high degree nodes connect to other high degree nodes. Together, these measures tells us how nodes integrate with other nodes of similar or different degree across the network.

#### Efficiency measures

The networks capacity for efficient information processing was determined using the global, local and diffusion efficiency, and communicability for each network (Latora and Marchiori, 2001; Goñi et al., 2013; Lella and Estrada, 2020). Global and local efficiency describes how easily information is transmitted across the network (globally) and around a given node (locally around a local subgraph given by a node and it’s neighbors), while also informing us of the fault tolerance of the network. Diffusion efficiency is given by the inverse of the mean first passage time, which denotes the expected number of steps for a random walker to pass from one node to another given node in the network. Similarly, communicability offers an alternative measures to the shortest path of the network by taking into account all shortest paths between two nodes (Estrada et al., 2012; Lella and Estrada, 2020).

Normalization was performed by dividing each communication value for a network with the mean of the same value found for a set of random networks. Random networks were generated such that they were equivalent to the network to be compared with (i.e. same number of nodes and edges). A total of ten random networks were used as the basis for normalization of each network.

### Software and tools

All simulations and subsequent analysis was performed using the Python programming language (Python Software Foundation, https://www.python.org/). Network parameters were analyzed and determined using the Python packages NetworkX, CDlib and netneurotools (Hagberg et al., 2008; Rossetti et al., 2019; Liu et al., 2025). Data was processed using the Python package Pandas (Wes McKinney, 2010) and figures were generated using the Python packages Matplotlib and Seaborn (Hunter, 2007; Waskom, 2021). Large language models (Microsoft Copilot, Google Gemini, GitHub Copilot) were used to generate parts of the code used for analysis and visualization. All generated code was reviewed before application, and unit testing was performed where applicable.

## AUTHOR CONTRIBUTIONS

NC: Conceptualization, Methodology, Software, Investigation, Formal Analysis, Visualization, Writing - Original Draft, Review and Editing. IS and AS: Conceptualization, Funding Acquisition, Project Administration, Supervision, and Writing — Review and Editing.

## DATA AVAILABILITY

The source code to generate network models can be found on GitHub:spatial-network-modeling.

## Supplementary Note A: Probability density functions

Dependence on the distance for the probability density functions of a connection occurring (Fig. A.1). The distance used is the distance in the range [0.001, 1]. Both probabilities for making connections are large for *d <* 0.01. Beyond this the log-normal connection probability decreases fast but maintains a heavy-tail whereas the exponential probability shows a preference for intermediate ranges up to *d <* 0.2, beyond which the probability declines fast.

The values used for *α* and *σ* in both connection probabilities with the corresponding mean degree can be found in Table 1. Each connection probability was tuned to have the absolute value of 50 = ⟨*k*⟩ was as close to zero as possible for the given spatial configuration. Each value was optimized over a range containing the the value of interest, and a mean of 5 networks were used to optimize ⟨*k*⟩ .

**Fig. A.1.**
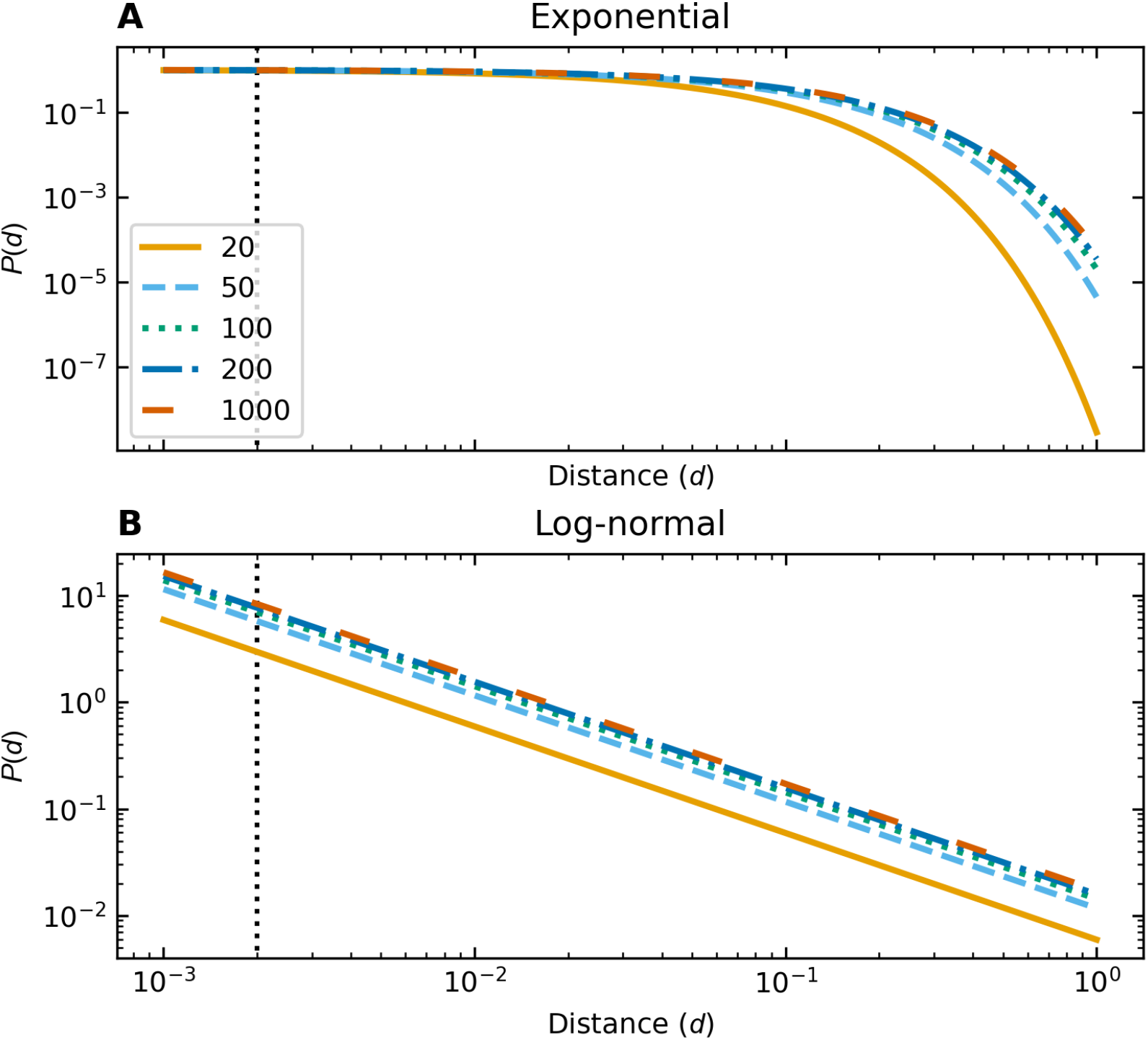
Probability density function *P* (*d*) of a connection being made between two nodes as a function of the distance *d* for different number initially placed nodes (20, 50, 100, 200 or 1000). **(A)** Probability density function *P* (*d*) for the exponential connection probability. **(B)** Probability density function *P* (*d*) for the log-normal connection probabilities. The vertical dotted line indicates the shortest allowed distance between two nodes in the systems. For the exponential probability *β* = 1, while *µ* = 0 in the log-normal case. For both connection probabilities *α* and *σ* were chosen such that the mean degree ⟨*k*⟩ of the network was ⟨*k*⟩ ≈ 50, resulting in slightly different profiles for different spatial distributions of nodes. Legends are shared between figures.

## Supplementary Note B: Raw and relative network parameters

The global, local and diffusion efficiency, and communicability were estimated for all networks both for increasing number of initially placed nodes *N*_init_ Fig. B.1 and for *N*_init_ = 20 with increasing pruning of long-range connections Fig. B.2. The raw data shows the data without any transformations while the relative data was determined according to Goñi et al. (2013).

For networks undergoing pruning, the pruning resulted in disconnected networks. These were excluded in any analysis. For the networks generated using the exponential wiring probability, most pruning densities resulted in disjoint networks Table 2. As such, all of these networks were excluded from the analysis and only networks generated using the long-range log-normal wiring probability were analyzed.

**Table 2.** The number of connected and unconnected networks for each wiring probability at all pruning degrees.

| Pruning Degree | Exponential |  | Log-normal |  |
| --- | --- | --- | --- | --- |
|  | False | True | False | True |
| 0.0 | 0 | 100 | 0 | 100 |
| 0.005 | 64 | 36 | 0 | 100 |
| 0.01 | 84 | 16 | 0 | 100 |
| 0.05 | 100 | 0 | 8 | 92 |
| 0.1 | 100 | 0 | 21 | 79 |

**Fig. B.1.**
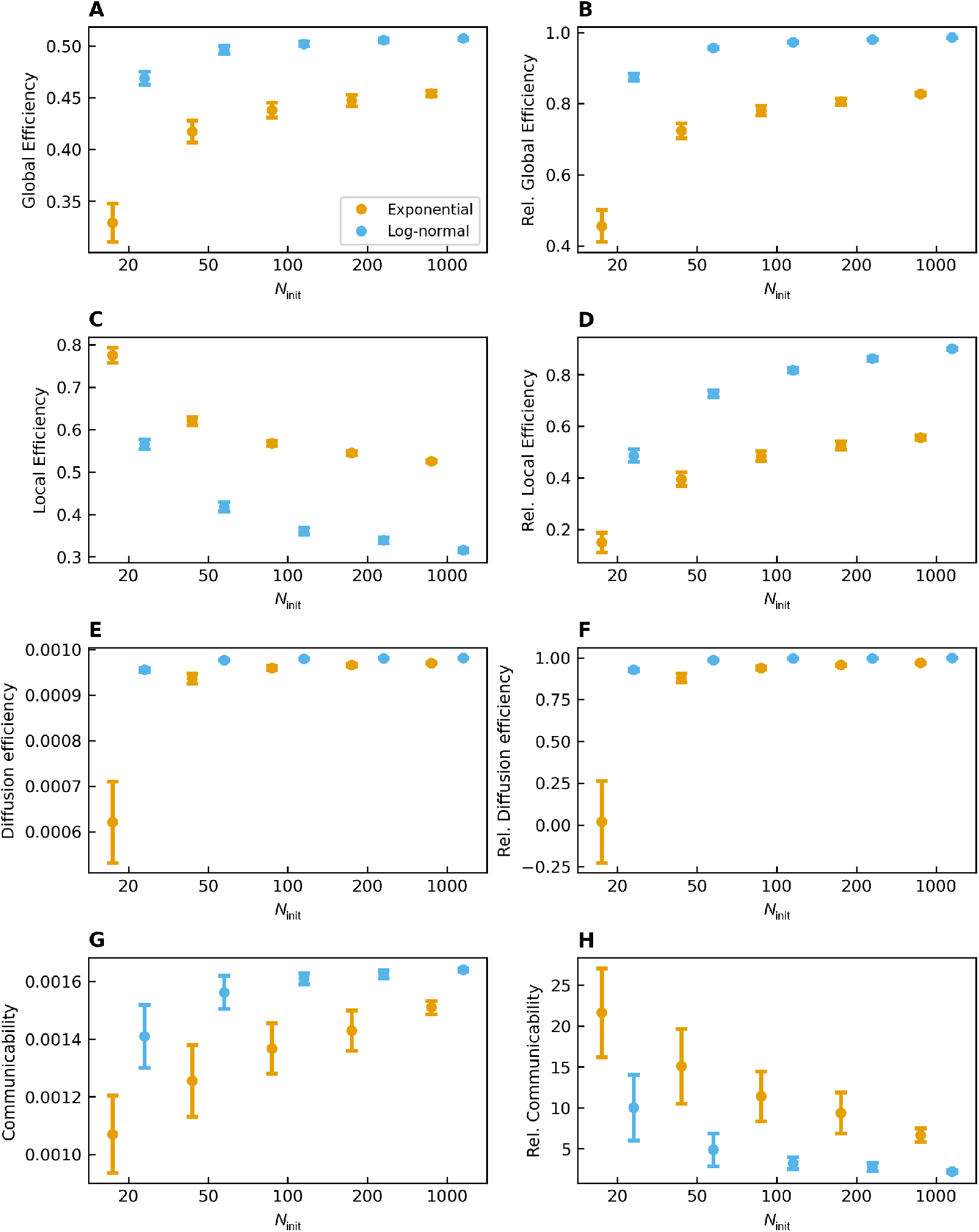
Raw and relative values for efficiency measures for all networks tuned to have a mean degree of ⟨*k*⟩ = 50 and increasing initially placed nodes *N*_init_. **(A)** The raw and **(B)** relative global efficiency, **(c)** raw and **(D)** relative local efficiency, **(E)** raw and **(F)** relative diffusion efficiency, and **(G)** raw and **(H)** relative communicability. The log-normal wiring probability (blue) show better capabilities related to global communication, whereas he exponential wiring probability (orange) show better local communication for all spatial conformations. For global efficiency and diffusion efficiency the relative and relative values follow the same trend, while the trend is inverted for local efficiency and communicability. All systems have *N* = 1000 nodes, and 100 instances were generated for each system. Each figure shows the mean generated from all realizations of each configuration with error bars showing the standard deviation. Legends are shared between figures.

**Fig. B.2.**
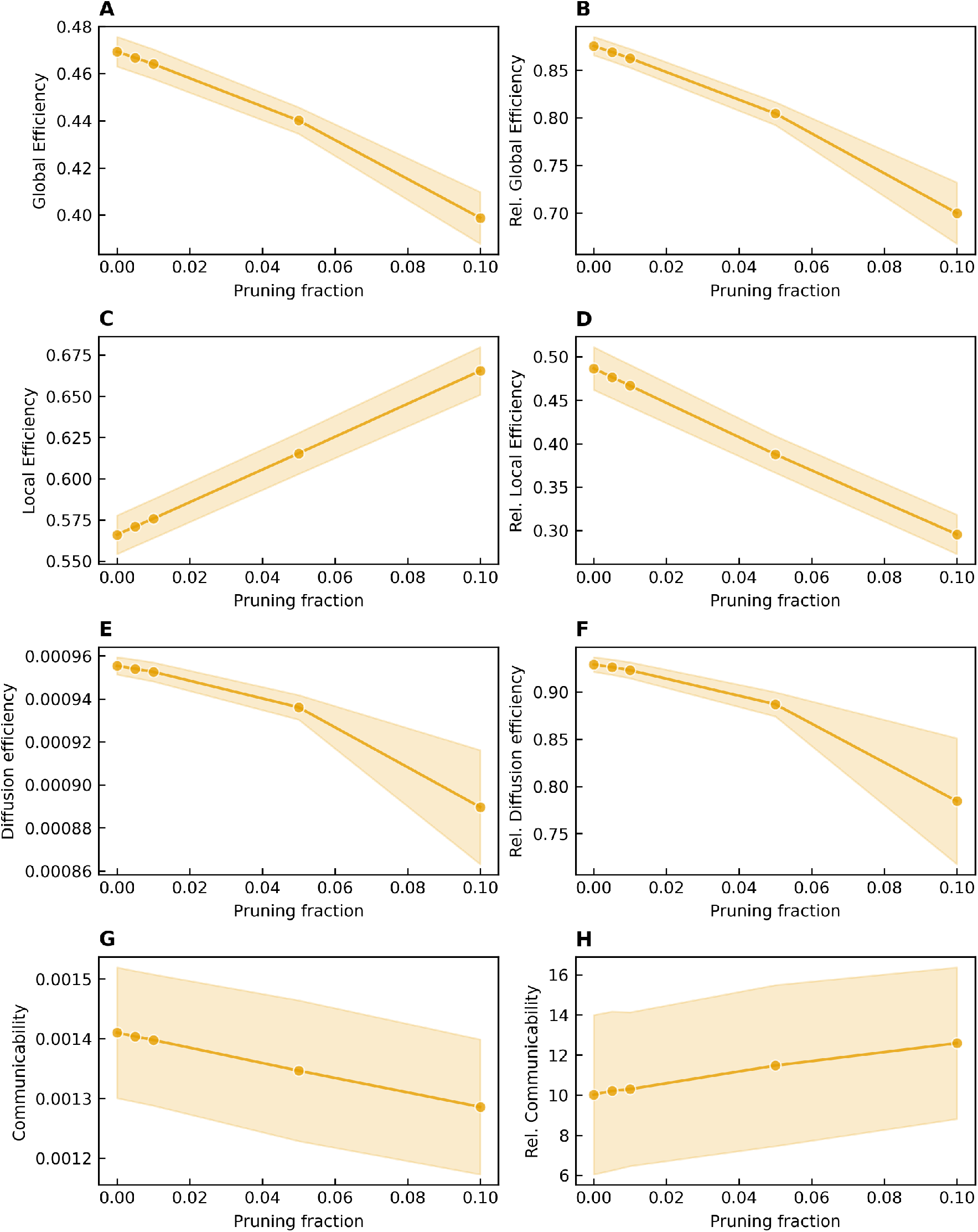
Raw and relative values for efficiency measures for networks generated using the log-normal wiring probability and initial nodes *N*_init_ = 20 at varying pruning degrees of long range connections. **(A)** The raw and **(B)** relative global efficiency, **(c)** raw and **(D)** relative local efficiency, **(E)** raw and **(F)** relative diffusion efficiency, and **(G)** raw and **(H)** relative communicability. Global efficiency and diffusion efficiency show a decreasing trend for raw and relative values. Local efficiency and communicability show opposite trends for the raw and relative values, with the raw data increasing and decreasing for local efficiency and communicability, respectively. All systems have *N* = 1000 nodes and the same networks were pruned at varying degree. Only fully connected networks were analyzed leading to a slight reduction in the number of networks for the strongest pruning degrees (92 networks at a pruning degree of 0.05 and 79 networks at a pruning degree of 0.1.). Each figure shows the mean generated from all realizations of each configuration with the shaded area showing the standard deviation

